# *Nicotiana tabacum* and *Vitis vinifera* as suitable growth-chamber models for microinjection-based studies of *Xylella fastidiosa*

**DOI:** 10.64898/2026.09.18.752583

**Authors:** David Labarga, Ana Cuevas, Miguel López-Belmonte, Natalia García-Tomsig, Marta Robledo

## Abstract

*Xylella fastidiosa* is one of the most important plant pathogens worldwide, infecting more than 700 plant species and causing major economic losses in affected agricultural regions. In the absence effective curative treatments under field conditions, experimental research is essential. However, research efforts are severely constrained by strict phytosanitary regulations and the pathogen’s restricted geographic distribution, as well as its demanding culture requirements. To the best of our knowledge, no studies have compared different host infection models under controlled growth-chamber conditions.

To address these limitations, we developed and compared a microinjection-based inoculation protocol across herbaceous and woody hosts maintained in biosafety growth chambers. Stems of *Vitis vinifera*, *Nicotiana tabacum*, *N. benthamiana*, and *Medicago sativa* were inoculated with *X. fastidiosa* by microinjection and maintained under controlled growth-chamber conditions. Pathogen establishment was assessed by qPCR analysis of petiole tissue, together with disease progression, plant growth, and leaf scorch development. Approximately one month after inoculation, bacterial detection rates reached 100% in *V. vinifera* and 80% in *N. tabacum*. Both species subsequently exhibited measurable disease development, whereas bacterial establishment and phenotypic responses were less consistent in *N. benthamiana* and *M. sativa*.

These results identify *N. tabacum* as a practical herbaceous model for preliminary screening and *V. vinifera* as a relevant woody host for subsequent validation. Their complementary use provides a controlled experimental framework for prioritizing candidate treatments, virulence factors, and resistance-related traits before evaluation under greenhouse or field conditions.

## Introduction

*Xylella fastidiosa* is a xylem-restricted bacterium with an exceptionally broad host range, capable of infecting more than 700 plant species worldwide (Cavalieri *et al*., 2025; Mourou *et al*., 2025). The pathogen can spread over long distances through the anthropogenic movement of plant materials and vectors (Morelli *et al*., 2021), and reports of *X. fastidiosa* outbreaks have increased markedly over the past decade (Landa *et al*., 2022). The economic impact is substantial, with estimated annual production losses across the EU reaching approximately €7.1 billion under a full-spread scenario, risking more than 540 000 jobs, according to a 2025 update from the European Commission’s Joint Research Centre (Joint Research Center, 2025). *X. fastidiosa* is regulated as a quarantine organism under EU Regulation 2016/2031, and working with this pathogen requires specialized containment facilities, substantially increasing research costs and limiting experimental capacity (Mourou *et al*., 2025).

Multiple strategies have been investigated to limit pathogen spread and reduce disease impact. First, regulatory containment and eradication measures based on surveillance and the establishment of demarcated zones have been adopted across Europe (Kyrkou *et al*., 2018; Morelli *et al*., 2021; Mourou *et al*., 2025). Second, vector control strategies employing insecticides, vegetation management, and natural enemies have been widely explored (Kyrkou *et al*., 2018; Saponari *et al*., 2019; Picciotti *et al*., 2021; López- Mercadal *et al*., 2022; Portaccio *et al*., 2025). Third, plant breeding and the deployment of resistant or tolerant cultivars represent one of the most sustainable approaches under investigation (Saponari *et al*., 2019; Landa *et al*., 2022; Mourou *et al*., 2025). Fourth, direct bacterial control using chemical compounds such as minerals and antimicrobial peptides has been attempted (Bragard *et al*., 2019; Portaccio *et al*., 2025). Finally, biological-control approaches have investigated endophytes, antagonists, and bacteriophages (Kyrkou *et al*., 2018; Bragard *et al*., 2019; Landa *et al*., 2022; Mourou *et al*., 2025). However, no single measure has benn shown to eliminate the bacterium under field conditions, and effective, long-lasting management strategies remain limited (Mourou *et al*., 2025). In the absence of effective curative treatments, experimental research using controlled laboratory systems is essential to advance understanding of pathogen-host interactions and evaluating potential control measures.

(Rogers, 2012)(Rogers, 2012)Herbaceous models such as *Arabidopsis thaliana* have been used to investigate pathogenesis and host–pathogen interactions (Rogers, 2012), but most experiments have been conducted in greenhouse facilities or under field conditions rather than in controlled laboratory growth chambers, limiting their broader accessibility. *Medicago sativa* has been described to produce visible immune responses within four months, thereby enabling evaluation of virulence factors and transcriptomic responses (Kubaa *et al*., 2019). However, its use as an experimental host remains limited. Inoculation studies in *Nicotiana* have been performed in biosafety greenhouses (Lopes *et al*., 2020; Pereira *et al*., 2020; Buisac *et al*., 2026), and nanoparticle-based treatments have been evaluated using soil drench applications in potted plants (Shantharaj *et al*., 2023). To the best of our knowledge, only two previous studies have evaluated *X. fastidiosa* infection under controlled growth-chamber conditions, both using *N. tabacum* (Francis et al., 2008; Tatulli et al., 2024), and no study has comparatively evaluated multiple host species.

Experimental research becomes even more complex in woody hosts, particularly in agronomically relevant species such as olive and grapevine. Olive is a primary agronomically relevant host, with experimental validation typically conducted under controlled greenhouse conditions using potted plants, though inoculation to symptom development requires periods exceeding one year (Saponari *et al*., 2017, 2019). (Hopkins, 2005; Kahn et al., 2023; Perelló et al., 2025)(Hopkins, 2005; Kahn et al., 2023; Perelló et al., 2025)Similarly, in vivo evaluations in grapevine, one of the most socioeconomically important crops globally (van Leeuwen *et al*., 2024), have been largely conducted in certified quarantine greenhouses or in affected regions. Field experiments are consequently concentrated in areas where the pathogen is established, including the Balearic Islands and the United States, whereas research in pest-free territories is restricted by phytosanitary regulations and limited access to high-containment facilities (Hopkins, 2005; Yang *et al*., 2011; Das *et al*., 2015; Cornara *et al*., 2016; Nascimento *et al*., 2016; Hao *et al*., 2017; Baccari *et al*., 2019; Zhang *et al*., 2019; Kahn *et al*., 2023; Martínez *et al*., 2023; Burbank *et al*., 2024; Wallis & Gorman, 2024; Perelló *et al*., 2025).

These geographic and infrastructural constraints highlight the need for controlled and reproducible infection protocols that can be applied comparatively across host species. Therefore, we developed and evaluated a microinjection-based inoculation methodology for *X. fastidiosa* in multiple plant hosts, including herbaceous (*Medicago sativa*, *Nicotiana benthamiana*, *Nicotiana tabacum*) and woody (*Vitis vinifera*) species, conducted entirely under controlled laboratory growth chamber conditions. Our objectives were to: (i) evaluate the performance of the microinjection protocol across different plant hosts; (ii) characterize bacterial colonisation dynamics and disease progression in each species; and (iii) identify suitable plant model system for future *X. fastidiosa* laboratory research and management strategy evaluation.

## Materials and Methods

### Plant material and growth conditions

Seeds of *Medicago sativa cv.* Aragón were surface-sterilized with 3.5% (v/v) sodium hypochlorite for 1 min, thoroughly rinsed with sterile distilled water (SDW), and imbibed at 4 °C for 5-7 h in darkness. Imbibed seeds were placed on water-agar plates (15 g L^-1^) and incubated at 24 °C in the dark for 24 h to induce germination. Uniformly germinated seedlings were transplanted into 200 mL (7 x 7 x 6) pots containing a 1:1 (v/v) mixture of universal substrate and vermiculite and transferred to a growth chamber maintained at 24/20 °C (light/dark) under a 16/8 h photoperiod.

*Nicotiana benthamiana* plants were established by surface-sowing 4-5 seeds per cell in 3 x 3 cm trays containing water-saturated soil substrate. The trays were maintained in transparent humidity chambers (≥60% RH) in a growth cabinet set to 24-26 °C and a 16/8 h (light/dark) photoperiod. Seedlings displaying 1-2 true leaves were transplanted under high humidity. When plants reached approximately 7 cm in height, they were transferred to 75-mL pots (5 cm diameter), containing the same soil substrate. Fertilization was supplied as a single top-dress application of a controlled-release NPK fertilizer (15:8:16; Premium Cote) at 5 g per pot, lightly incorporated into the soil prior to thorough watering.

Seeds of *Nicotiana tabacum* cv. Havana (Magic Garden Seeds, Germany) were sown in seedling trays filled with a 3:1 mixture of universal substrate and vermiculite, moistened to saturation. Trays were sealed with a transparent lid to maintain high humidity and were placed in a climate chamber under a 16/8-hour light/dark photoperiod at 24-26 °C (light) and 19 °C (dark). Individual seeds were positioned on the substrate surface. Germination occurred within 7-10 days, with >90% germination success. Seedlings reaching 2-3 cm in height (∼3-4 weeks) were transplanted to 7x7x6 pots (200ml) using the same substrate mixture. Following transplantation, grown in a growth chamber under the same germination conditions until they reached sufficient size for inoculation experiments.

*Vitis vinifera* L. cv. Tempranillo clusters, clone RJ43, were harvested in September 2025. Seeds were manually extracted from the berries and selected by flotation in water, discarding floating seeds and retaining sinking seeds for subsequent treatments. Prior to germination, seeds were stored at 4 °C with Cu(OH)_2_ 94% (ThermoScientific). Germination was performed following Rodríguez (2019). Briefly, seeds were then scarified by immersion in 10% H_2_SO_4_ (Sigma-Aldrich, USA) for 1.5 min under vortex agitation, rinsed thoroughly with water, and kept under these hydration conditions for 24 h. Seeds were subsequently placed in cloth bags and stored in vermiculite at 4 °C until germination. For germination, cloth bags containing the seeds were transferred to moistened vermiculite and subjected sequentially to a constant regime of 35 °C and 90% relative humidity (RH) for 48 h in darkness, followed by an alternating regime of 32 °C for 8 h and 22 °C for 16 h at 90% RH in darkness for 10 days, and finally a 16 h photoperiod (125 µmol m⁻² s⁻¹) at 25 °C combined with 8 h darkness at 22 °C, at 80% RH, maintained until transplantation. Thirty-five seedlings were transplanted into pots (7 x 7 x 8 cm) with a mixture of peat and vermicompost (8:1).

Throughout the experimental period, all plants were watered on demand to maintain adequate substrate moisture and prevent water stress.

### *Xylella fastidiosa* culture and plant microinjection

*X. fastidiosa* subsp. *fastidiosa* (Xff) strain Tem1 was used to inoculate in *N. tabacum* and *V. vinifera* whereas Xff strain 5235 was used for *M. sativa* and *N. bethamiana*. In addition, *M. sativa* was inoculated with *X. fastidiosa* subsp. *pauca* strain DeDonno (Xfp)*, fastidiosa* and *multiplex* strain 5901 (Xfm). The strains were retrieved from cryogenic storage and cultured on solid Pierce’s Disease 3 (PD3) medium(Davis *et al*., 1981). After 7 to 10 days of incubation at 28 °C, bacterial colonies were subcultured under identical conditions to obtain fresh bacterial growth for inoculation. For plant inoculation, mucoid colonies were harvested from the agar surface and suspended in sterile liquid PD3 medium. The concentration of the bacterial suspension was adjusted to an optical density corresponding to 10^7^-10^8^ CFU ml^-1^, as verified by serial dilution and plate counting.

Plants were microinjected with a total volume of 10 µL of the bacterial suspension, split into two separate 5 µL injections performed at two distinct areas within the same internode near the soil level. The suspension was delivered through a 25-gauge needle using a syringe pump (Model NE-1000; New Era Pump Systems Inc., USA). At the time of inoculation, the age of the host plants was as follows: *Vitis vinifera*, 16 weeks old; *Nicotiana benthamiana*, 10 weeks old; *Nicotiana tabacum*, 6 weeks old; and *Medicago sativa*, 12 weeks old. Control plants were microinjected with the same volume of sterile water.

### Plant growth and symptom assessment

Stem length was measured weekly for all plants from inoculation until the end of the experiment using a ruler to the nearest millimeter.

Leaf scorch symptoms were periodically assessed after inoculation until symptom stabilization or plant senescence. Assessment methods differed among species because of practical constraints and differences in plant architecture.

For *M. sativa*, the total number of scorched leaves were recorded for all plants within each treatment at each day post-inoculation (DPI). This cumulative value was divided by the number of plant replicates within the corresponding treatment to the mean number of scorched leaves per plant at each time point. Because symptoms were recorded collectively rather than for individual plants, replicate-level data were unavailable, precluding inferential statistical comparisons between treatments.

For *Nicotiana benthamiana*, *N*. *tabacum*, and *Vitis vinifera*, leaf scorch was assessd for each plant individually. At each observation time, the number of leaves displaying marginal necrosis or scorch and the total number of leaves were recorded. The percentage of scorched leaves per plant was calculated as:

Scorched leaves (%) = (number of scorched leaves / total number of leaves) x 100.

These individual plant-levelvalues were used for subsequent statistical analysis.

### Standardization of disease progression among host plants

Disease progression was quantified by calculating the Area Under the Disease Progress Curve (ADPC) according to the trapezoidal integration method described by Fry (1978). Briefly, this parameter integrates both the onset and the rate of epidemic development into a single value by accumulating the proportion of affected plant recorded at successive intervals throughout the evaluation period.

To account for variations in disease duration and allow for direct comparison across different experimental timelines, the standard ADPC values (expressed in days- proportion) were normalized to obtain the relative ADPC (rAUPCD). This standardization was achieved by dividing the calculated ADPC by the maximum theoretical area of the graph, calculated as the duration from inoculation to the final observation multiplied by the maximum disease severity score used for the corresponding host. To this end, we established a disease severity scale for each plant species, as shown in Fig. 1. Based on each classification, we assigned a score to the damage observed on each plant studied.

**Fig. 1.**
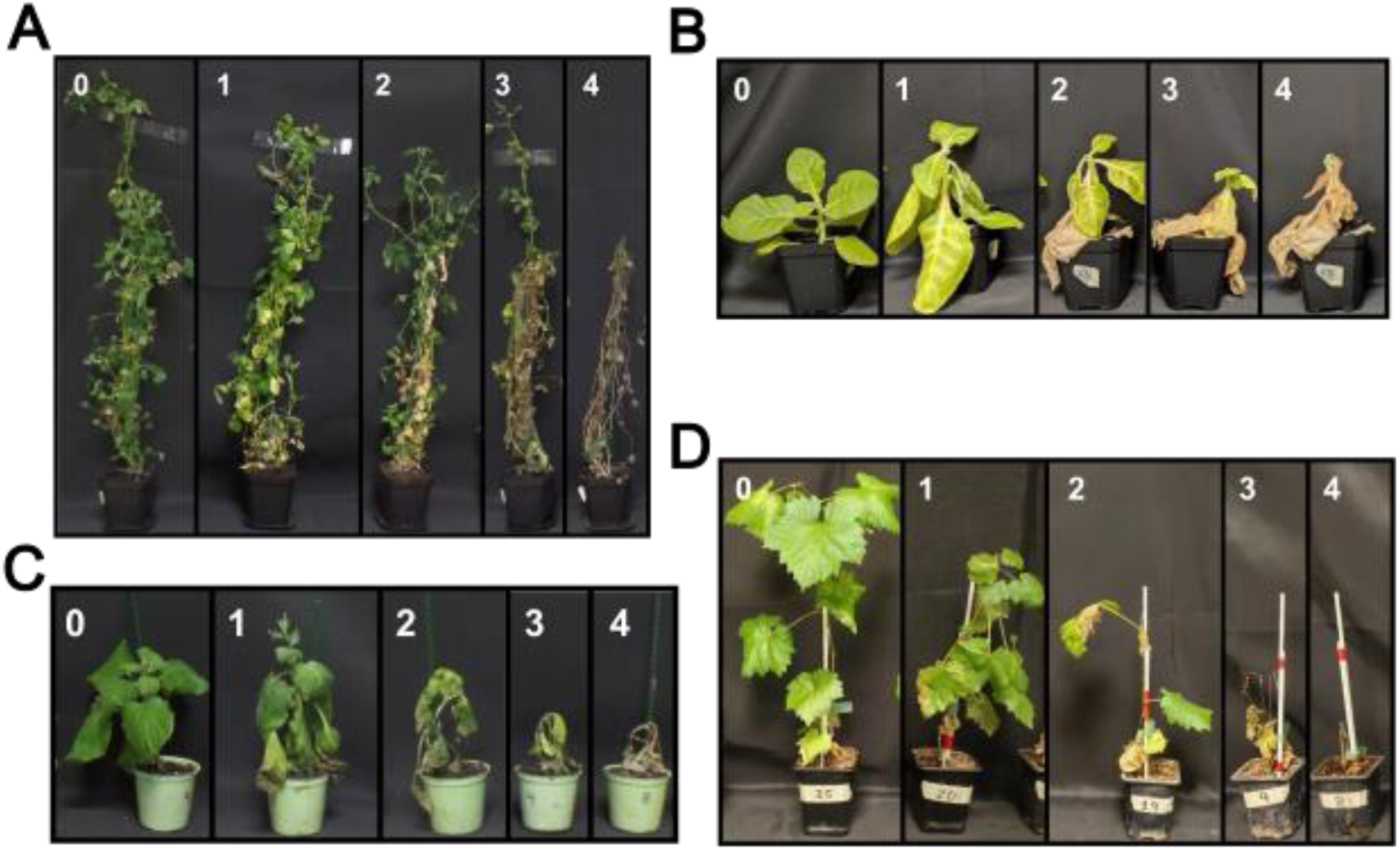
Disease severity scales used to assess symtoms associated with *Xylella fastidiosa* inoculation in *Medicago sativa* (A), *Nicotiana tabacum* (B), *Nicotiana benthamiana* (C) and *Vitis vinifera* (D). Symptoms were rated from 0 to 4: 0, no symptoms; 1, mild or nonspecific symptoms; 2, clear symptoms, including leaves with significant marginal necrosis; 3, severe symptoms affecting a high percentage of leaves; 4, plant death.

### DNA extraction and qPCR

To assess *Xylella* infection progression, DNA was extracted from the plants petioles. Specifically, 100 mg of petiole tissue was collected from the leaf located immediately above the inoculated internode. DNA extraction was performed using the NucleoSpin® Plant II kit (Macherey-Nagel, Düren, Germany) according to the manufacturer’s instructions with a final elution volume of 100 µL.

Detection of *X. fastidiosa* was performed using the real-time PCR assay described by Harper et al. (2010), targeting the *rimM* gene. All samples were analyzed in three technical replicates. Each qPCR plate also included three negative controls and three no-template controls (NTCs), in which the DNA template was replaced with sterile distilled water. The primers were XF-F (5′- CACGGCTGGTAACGGAAGA -3′) and XF-R (5′- GGGTTGCGTGGTGAAATCAAG-3′), and the hydrolysis probe XF-P (5′- FAM- TCGCATCCCGTGGCTCAGTCC-BHQ1-3′). Reactions were performed in a final volume of 20 µL mixture containing 2 µL of DNA template, 4 µL of PerfeCTa qPCR ToughMix (Quantabio, Beverly, MA, USA), 0.6 µL of each primer (10 µM), 0.2 µL of probe (10 µM), and 12.6 µL of sterile distilled water. Amplification was conducted on an Applied Biosystems^TM^ StepOnePlus Real-Time PCR Systems (Thermo Fisher Scientific). The termal cycling programme consisted of an initial denaturation step of 10 min s at 95 °C, followed by 40 cycles of 95 °C for 10 s and 62 °C for 40 s.

To establish a quantitative relationship between qPCR cycle threshold (Ct) values and bacterial load, a matrix-matched standard curve was constructed following the general principles described by Baró et al. (2020, 2021), with modifications. Briefly, 100 mg of *Medicago sativa* petioles was mixed with 100 µL of bacterial suspension (OD_600_ = 6) at serial dilutions. The viable bacterial concentration (CFU ml⁻¹) for each dilution was independently determined by plating on solid PD3 medium. DNA extraction from the mixed plant tissue and qPCR were performed following the procedures described above. The standard curve (Fig. S. 1) was constructed by plotting mean Ct values against the logarithmic values of initial bacterial concentrations (CFU ml⁻¹). Linear regression analysis yielded a predictive equation that allowed the conversion of Ct values obtained from experimental samples into quantitative bacterial load estimates.

Estimated bacterial loads were derived from the matrix-matched calibration curve and expressed as CFU equivalents per gram of petiole tissue. For construction of the calibration curve, 100 µL (0.1 mL) of each bacterial dilution was added to 100 mg (0.1 g) of plant tissue. Therefore, the numerical bacterial concentration of the initial suspension expressed as CFU mL^-1^ was equivalent to the bacterial load introduced into the plant matrix expressed as CFU g^-1^ (e.g., 10^6^ CFU mL^-1^ x 0.1 mL / 0.1 g = 10^6^ CFU g^-1^). The threshold for classifying samples as positive was established based on the bacterial-load estimates obtained from non-inoculated plant controls. Samples yielding values within this background range were considered negative.

### Statistical analysis

All analyses were conducted in R version 4.5.1 using the packages glmmTMB (version 1.1.13) for mixed-effects modeling (using always plant replicate as a random intercept in longitudinal analyses. Type II Wald χ² tests were performed using the Anova function in the car package. Post hoc comparisons were conducted using emmeans, with Tukey adjustment for multiple comparisons. Candidate response distributions were explored using fitdistrplus, and model fit was assessed. All the models were adjusted to their best distribution. Significance was set at α = 0.05 for all statistical tests.

Stem length measurements were modeled as continuous response variables using generalized linear mixed models (GLMM). For the combined-species analysis, models included species, treatment, DPI, and their relevant interactions as fixed effects, with plant identity as random effect. Linear, quadratic, and cubic of time were compared using likelihood-ratio tests to evaluate whether growth trajectories exhibited nonlinear patterns. Separate models were subsequently fitted for each species, and polynomial complexity was selected independently for each dataset.

Leaf scorch data were analyzed separately for each species due to differing patterns of symptom expression. For *N. benthamiana*, binomial GLMM models were fitted at the individual plant level. Early time points presenting structural zeros and complete separation were excluded from inferential models to ensure mathematical convergence because the complete absence of symptoms in the control group resulted in zero variance and complete separation. For *N. tabacum* and *Vitis vinifera*, GLMM with binomial family were fitted to model the probability of leaf scorch. The fixed effects structure was Treatment x DPI (as fixed factors).

Symptom severity was quantified over time using rAUDPC. Distribution selection via fitdistrplus identified beta distribution as optimal, reflecting the bounded (0, 1) nature of the rAUDPC response. Given the presence of zero values, we fitted a GLMM with ordinal beta family (ordbeta in packages glmmTMB) with Treatment x Species as fixed effects.

Type II Wald χ² tests were performed on the best model of each parameter to assess main effects and interactions. Post-hoc pairwise comparisons within each species were conducted using emmeans with Tukey adjustment for multiple comparisons at each observed timepoint (DPI).

## Results

### Matrix-matched Ct-to-CFU Regression Model for *X. fastidiosa* infection quantification (CFU ml^-1^) from qPCR-Ct values

Previous qPCR-based quantification approaches for *X. fastidiosa* differed in whether a plant matrix was included in the calibration. Baró et al. (2020) built standard curves solely from bacterial suspensions, without plant material, whereas Baró et al. (2021) used matrix-matched curves prepared from almond sap or tissue extracts fortified with known bacterial concentrations. Following this latter approach, a matrix-matched standard curve was constructed here by directly mixing known bacterial concentrations with homogenized *M. sativa* petiole tissue, having a Ct to CFU conversion.

The calibration curve encompassed serial dilutions of *X. fastidiosa* subsp. *fastidiosa*, (Xff) ranging from 2.12 to 8.97 log CFU ml^-1^, as determined by plate counting. The relationship between Ct values and bacterial concentration displayed a strong negative linear correlation (Fig. S. 1). Ct values ranged from 14.61 to 35.24, spanning a 20.63 cycles. Linear regression analysis of Ct values plotted against log (CFU ml^-1^) yielded the following predictive equation: log (CFU ml^−1^) = -0.33Ct+13.46. The coefficient of determination (R²) was 0.97, indicating excellent explanatory power across the tested concentration range. This equation was used to convert Ct values from experimental samples into estimated bacterial loads, expressed as CFU equivalents per millilitre of tissue homogenate. Because uncertainty increases near the upper Ct limit, samples producing estimates within or close to the range observed in non-inoculated controls were not classified as positive.

### *Medicago sativa* shows limited *X. fastidiosa* colonization and no detectable disease development under growth-chamber conditions

*M. sativa* (alfalfa) was selected as an experimental host to evaluate its susceptibility to *X. fastidiosa* infection as a legume model system, following previous reports suggesting susceptibility to this pathogen under greenhouse conditions (Kubaa *et al*., 2019; López-Mercadal *et al*., 2021; Ghanbari *et al*., 2024). A comprehensive phenotypic assessment was conducted to compare its response to inoculation with different strains of *X. fastidiosa* belonging to subsp. *fastidiosa* (Xff), *multiplex* (Xfm) and *pauca* (Xfp).

Bacterial colonisation was assessed by qPCR to determine whether infection had been successfully established (Fig. 2A). Uninfected control plants consistently yielded background qPCR signals of approximately 2-3 log CFU g^-1^. *M. sativa* showed only limited evidence of bacterial colonization, with 30% plants inoculated with Xff or Xfm showing bacterial loads clearly above those of non-inoculated controls at 190 days post-inoculation (DPI), reaching approximately 6 log CFU g^-1^. No plants Xfp-inoculated showed detectable bacterial loads above the control range at any sample time.

**Fig. 2.**
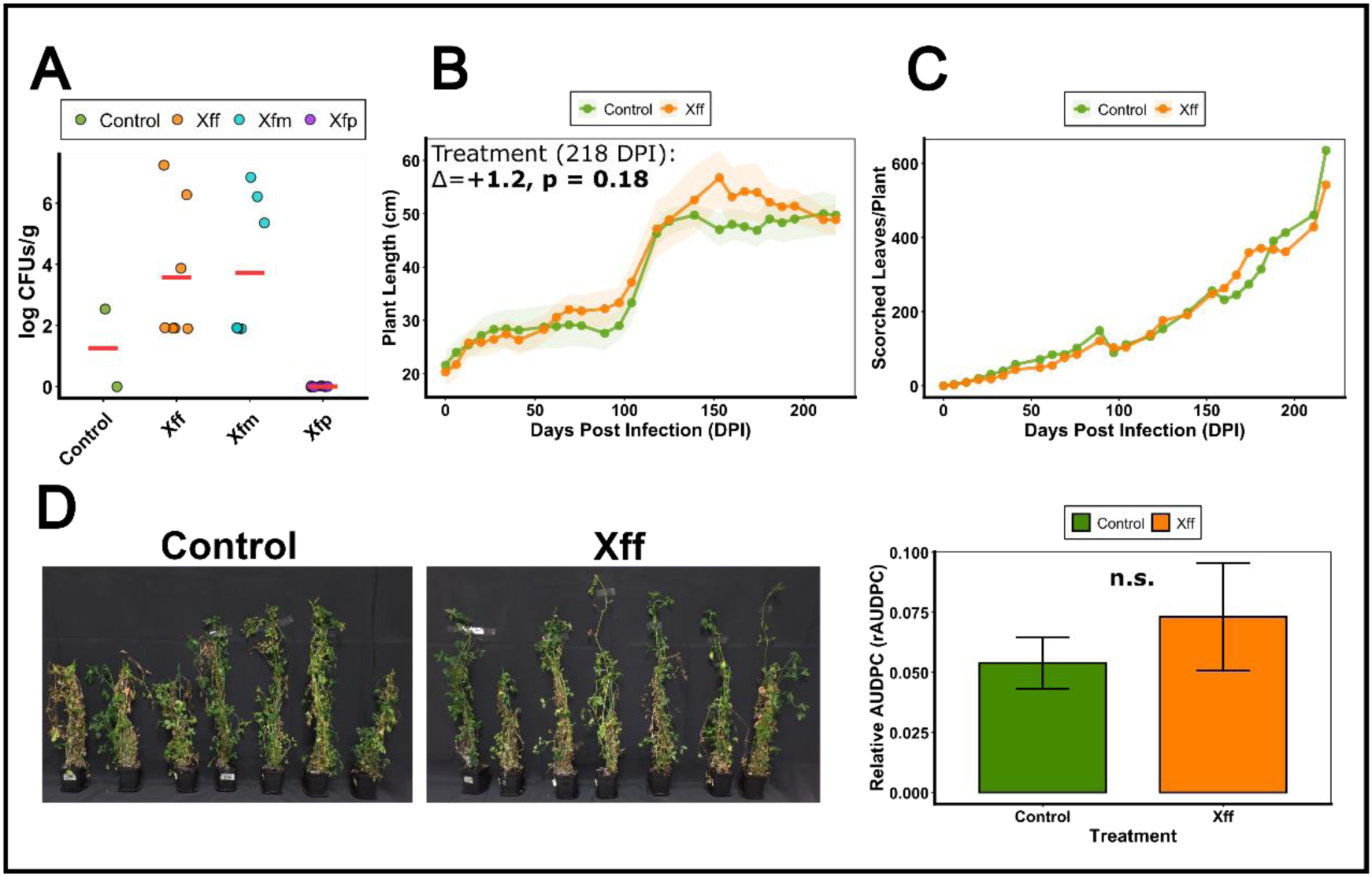
Assessment of *Xylella fastidiosa* colonization and disease progression in *Medicago sativa*. (A) Estimated bacterial titres in petiole tissues at 190 DPI. Each point represents one plant replicate. Red horizontal lines indicate the mean bacterial titre (log CFU g^-1^) for each treatment: non-inoculated controls (green), *X. fastidiosa* subsp. fastidiosa (Xff, orange), *X. fastidiosa* subsp. multiplex (Xfm, blue), and *X. fastidiosa* subsp. pauca (Xfp, purple). (B) Plant growth dynamics over time following inoculation with Xff (orange) compared to non-inoculated controls (green). Shaded areas represent SEM. The treatment effect at the final evaluation date (DPI 218) was assessed using mixed-effects model. The estimated difference between treatments at the final evaluation time point (218 DPI) and the corresponding p-value are shown in the graph. (C) Leaf scorch development over time. Values represent the mean number of scorched leaves per plant replicate for control (green) and Xff-inoculated (orange) plants. (D) Pictures of non- inoculated control plants and plants inoculated with Xff 213 days post-inoculation (DPI) (left), alongside the Relative Area Under the Disease Progress Curve (rAUDPC) for symptom severity (right). Bars represent means ± standard errors (SEM); green and orange correspond to control and Xff-inoculated plants, respectively. Pairwise comparison between treatments was performed using estimated marginal means (Tukey- adjusted). Significance codes: ns: not significant.

Symptom development of Xff-inoculated plants was quantified using the relative area under the disease progress curve (rAUDPC) across the 218-day experimental period (Fig. 2D and Table S. 1). Inoculated and control plants displayed comparable rAUDPC distributions with overlapping confidence intervals, and no significant treatment effect was detected (p = 0.92). Similarly, plants inoculated with Xfm and Xfp were phenotypically indistinguishable from non-inoculated controls throughout the experiment. *M. sativa*, further indicating that none of the three strains induced detectable disease development under the conditions tested (Fig. S. 2).

To enable comparison with the controls and other host species, we analyzed stem elongation and leaf scorched in Xff-inoculated plants, despite the lack of visible disease manifestation. Plant lenght followed broadly similar trajectories in inoculated and control plants (Fig. 2B and Table S. 2), with Xff-inoculated and control plants reaching approximately 50 cm by 218 DPI. Although the treatment × DPI interaction was significant (p = 0.02), the overall treatment effect and the final comparison were not significant (p = 0.54. Final stem lengths differed by only 1.2 cm (Δ = +1.2 cm, p = 0.19). Similarly, the proportion of scorched leaves increased over time in both treatments during the 218-day observation period (Fig. 1C and Table S. 3), but no significant differences were detected between inoculated and control plants.

Collectively, *M. sativa* showed limited bacterial establishment and no consistent phenotypic response following inoculation with three different *X. fastidiosa* stains. These results indicate that this species has limited suitability for experiments requiring reproducible systemic infection and disease development under controlled growth chamber conditions.

### *Nicotiana benthamiana* exhibits variable *Xylella fastidiosa* colonisation and phenotypic responses under growth-chamber conditions

Given the limited quantifiable responses observed in *M*. *sativa*, the experimental focus shifted to the genus *Nicotiana*, based on prior evidence of *X*. *fastidiosa* susceptibility in greenhouse assays (Baró *et al*., 2022). *N*. *benthamiana* was therefore selected to evaluate whether this species could pro as a more sensitive and informative model system for Xff infection under growth-chamber conditions.

**Fig. *3*Table S. *1*** Molecular assessment revealed variable colonisation success. Xff was clearly detected in 40% of the inoculated plants at 29 DPI (4/10), with bacterial titres ranging from 5 to 7 log CFU g^-1^ (Fig. 3A). Quantification of disease progression via rAUDPC revealed no significant differences between inoculated and control plants (p = 0.99) (Fig. 3D and Table S. 1). Vegetative growth was markedly affected by inoculation (Fig. 3B and Table S. 2). Control plants increased in height from 9 to 14 cm over the experimental period, whereas inoculated plants exhibited progressive growth impairment, with mean plant height decreasing from 15 cm at 15 DPI to 7 cm by 29 DPI. Although the overall treatment effect was not statistically significant (treatment effect: χ^2^ = 2.41, p=0.12), a significant treatment x DPI interaction was detected (χ^2^ = 12.8, p=0.005). At the final timepoint, inoculated plants were, on average, 7.56 cm shorter than controls (p < 0.05). No scorched leaves were observed through 14 DPI, preventing early-stage comparisons between treatments. However, by 29 DPI, the proportion of scorched leaves was significantly higher in inoculated plants than in controls (p < 0.05; Fig. 3C and Table S. 3), with a difference of 22.67% between treatments.

**Fig. 3.**
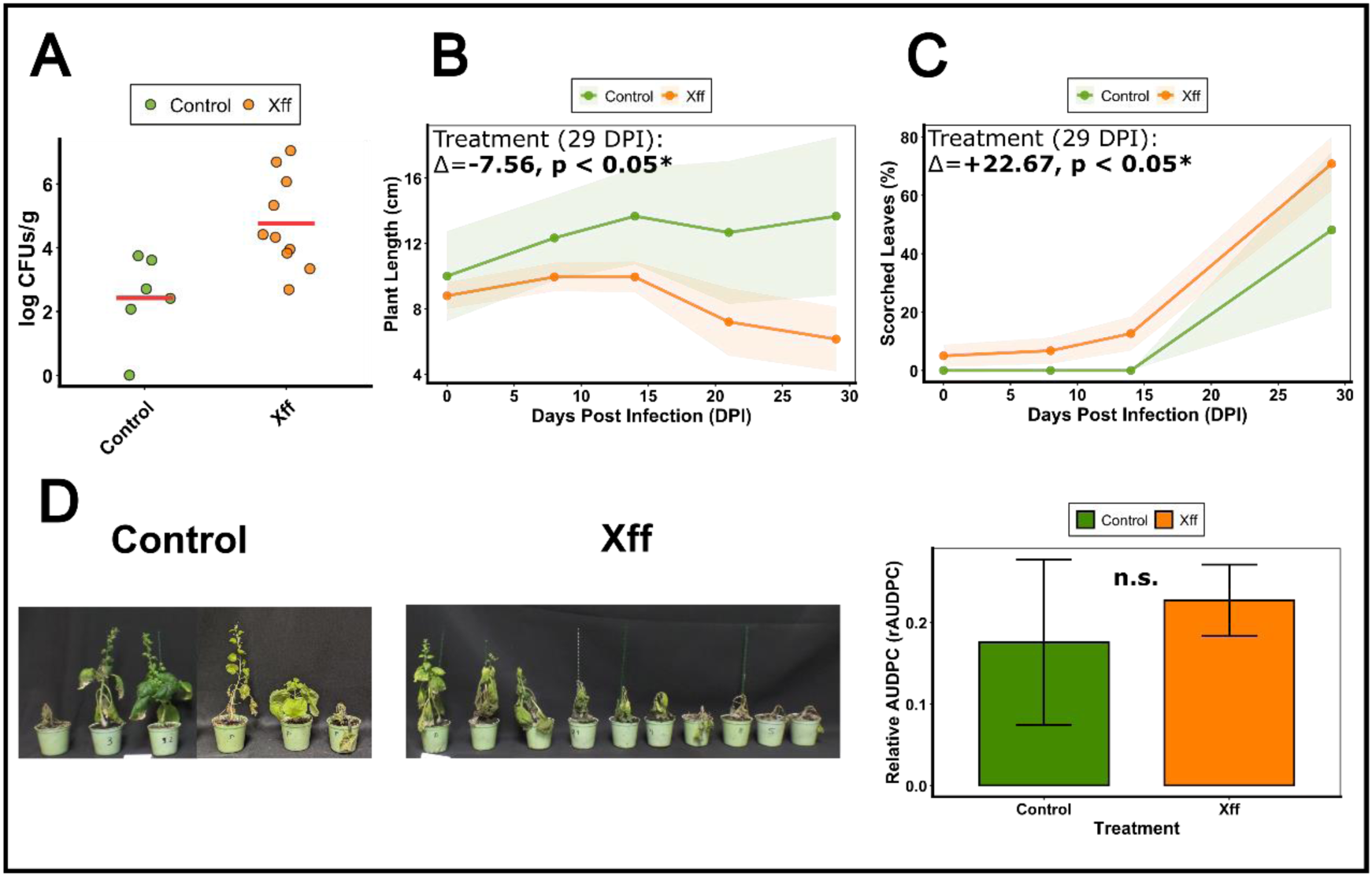
Assessment of *Xylella fastidiosa* infection and disease progression in *Nicotiana benthamiana*. (A) Estimated bacterial population levels in petiole tissues at 29 DPI. Each point represents an individual plant replicate. Red horizontal lines indicate the mean bacterial titres, expressed as log CFU g^-1^ value for each treatment: non-inoculated controls (green), *X. fastidiosa* subsp. *fastidiosa* (Xff, orange). (B) Plant growth dynamics over time following inoculation with Xff (orange) compared to non-inoculated controls (green). Shaded areas represent SEM. The effects of treatment, DPI, and their interaction were assessed using a two-way ANOVA. The difference between treatments at the final evaluation time point (29 DPI), together with the corresponding p-value, is shown in the graph. (C) Development of leaf scorch symptoms over time. Values represent the percentage of scorched leaves per plant replicate for control (green) and Xff-inoculated (orange) plants. (D) Representative non-inoculated control plants and plants inoculated with Xff at 213 days post-inoculation (DPI) (left), alongside the Relative Area Under the Disease Progress Curve (rAUDPC) for symptom severity (right). Bars represent means ± standard errors (SEM); green and orange indicate control and Xff-inoculated plants, respectively. Pairwise comparison between treatments was performed using estimated marginal means (Tukey-adjusted). Significance codes: ns: not significant, p > 0.05; *.

In contrast to *M. sativa*, *N. benthamiana* exhibited detectable phenotypic responses to *Xff* 5235 inoculation, particularly marked growth inhibition and delayed but detectable leaf scorch symptoms. Nevertheless, variable bacterial establishment and modest disease development limit its reproducibility as a growth-chamber infection model under the conditions tested.

### *Nicotiana tabacum* sustains high *X. fastidiosa* titres, marked growth suppression, and progressive disease development

*N. tabacum* has been reported to be susceptible to *X. fastidiosa* infection in multiple independent studies in greenhouses (Pereira *et al*., 2020; Shantharaj *et al*., 2023). This species was therefore incorporated into our experimental framework to assess whether it could prove more suitable experimental model than the previously tested hosts in growth chambers. The same comprehensive phenotypic assessment was applied to enable direct comparison of host responses across the experimental panel.

Bacterial establishment was frequent and sustained (Fig. 4A), with Xff detected in 80% of the inoculated plants at 35 DPI (8/10), 90% at 62 DPI (9/10), and 87.5% at 139 DPI (7/8; two plants died and was impossible to extracted the DNA). Bacterial titres were consistently high across the three sampling times, ranging from 6 to 8.2 log CFU g^-1^, indicating sustained systemic colonization throughout the experiment (Fig. 4A).

**Fig. 4.**
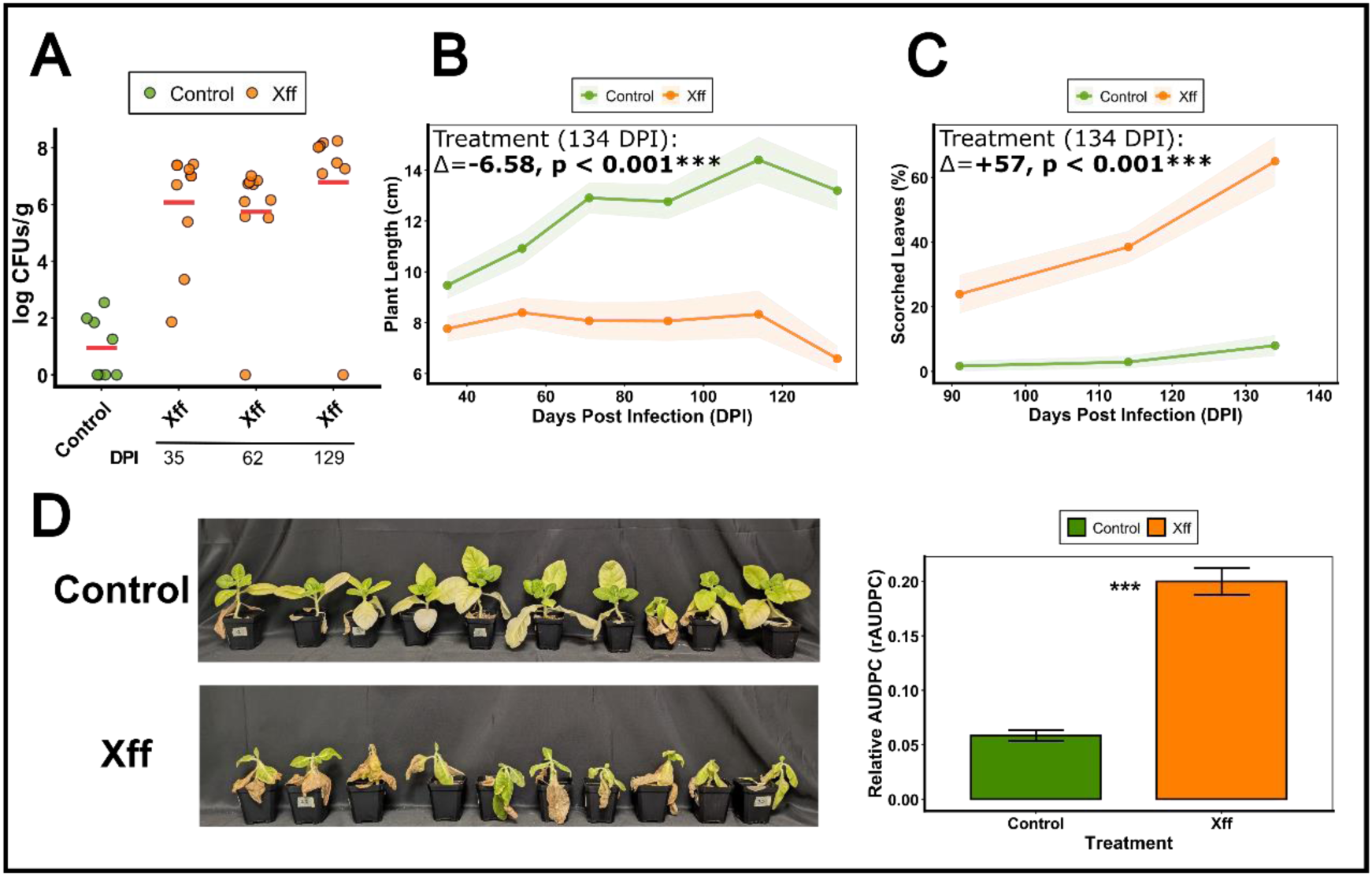
Assessment of *Xylella fastidiosa* infection and disease progression in *Nicotiana tabacum* in growth chambers. (A) Estimated bacterial population levels in petiole tissues at 35, 62, and 129 DPI. Each point represents an individual plant replicate. Red horizontal lines indicate the mean bacterial titre (log CFU g^-1^) value for each treatment: non-inoculated controls (green), and *X. fastidiosa* subsp. *fastidiosa* (Xff, orange). (B) Plant growth dynamics over time following inoculation with Xff (orange) compared to non-inoculated controls (green). Shaded areas indicate the SEM. The treatment effect at the final evaluation date (129 DPI) was assessed using a two-way ANOVA with treatment and DPI as fixed factors (Δ and p-value indicated in the graph). (C) Leaf scorch progression over time. Values represent the percentage of scorched leaves per plant. (D) Representative non-inoculated control plants and plants inoculated with Xff at 127 days post-inoculation (DPI) (left), alongside the Relative Area Under the Disease Progress Curve (rAUDPC) for symptom severity (right). Bars represent the mean ± standard error (SEM); green and orange correspond to control and Xff-inoculated plants, respectively. Pairwise comparison between treatments was performed using estimated marginal means (Tukey-adjusted). Significance codes: p ≤ 0.001; ***.

Disease progression in *N. tabacum* infected plants was also pronounced and clear. By 127 DPI, the control group keep a turgid, healthy green appearance, with only occasional lower-leaf shedding typical of the species’ natural turnover. Infected plants, however, suffered extensive foliar scorching and a marked systemic decline, culminating in plant death by this sampling date (Fig. 4D). rAUDPC values in inoculated plants reached approximately 0.20, significantly exceeding those of controls (≈0.05, p = 0.0002), indicating robust and progressive symptom accumulation (Fig. 4D and Table S. 1).

**Fig. *4***Stem elongation was severely restricted in inoculated plants (Fig. 4B and Table S. 2). Inoculated plants remained below 9 cm in height, whereas control plants reached approximately 14 cm by 120 DPI. Significant effects of treatment (χ^2^ = 37.04, p < 0.001), DPI χ^2^ = 32.25, p < 0.001), and their interaction (χ^2^ = 37.82, p < 0.001) were detected. At 134 DPI, inoculated plants were, on average, 6.58 cm shorter than control plants (p < 0.001).

Foliar symptoms differed significantly between treatments from DPI 90 onwards (Fig. 4C and Table S. 3), with inoculated plants displaying progressive and sustained symptom development throughout the observation period. Significant effects of treatment (χ^2^ = 39.13, p < 0.001), DPI (χ^2^ = 19.69, p < 0.001), and their interaction (χ^2^ = 27.87, p < 0.001) were detected. The greatest difference between treatments was observed at 134 DPI, when inoculated plants showed a 57-percentage-point increase in the proportion of scorched leaves relative to controls (Δ = +57, p < 0.001).

The Tem1-*N. tabacum* combination provided a highly reproducible infection system, exhibiting clear and consistent bacterial colonisation and phenotypic responses across all measured parameters. The sustained bacterial colonisation, progressive disease development, and marked plant growth suppression observed in this system support its use as a reliable model for studying *X. fastidiosa* pathogenesis under controlled growth-chamber conditions.

### *Vitis vinifera* supports consistent *X. fastidiosa* colonization and disease development under growth-chamber conditions

Following the robust infection outcomes observed in *Nicotiana tabacum*, the experimental framework was extended to a woody plant species. *Vitis vinifera* was selected because it is a naturally susceptible woody perennial of considerable economic importance that is severely affected by *X. fastidiosa* in agricultural systems worldwide (Landa *et al*., 2022; Mourou *et al*., 2025). The aim was to determine whether the controlled growth-chamber inoculation protocol could induce consistent bacterial colonization and disease development in this host.

Bacterial colonisation was consistent across inoculated plants (Fig. 5A), with all seven microinjected plants (100%) testing positive at 32 DPI, and reaching titres ranging from 4.6 to 7.7 log CFU g^-1^.

**Fig. 5.**
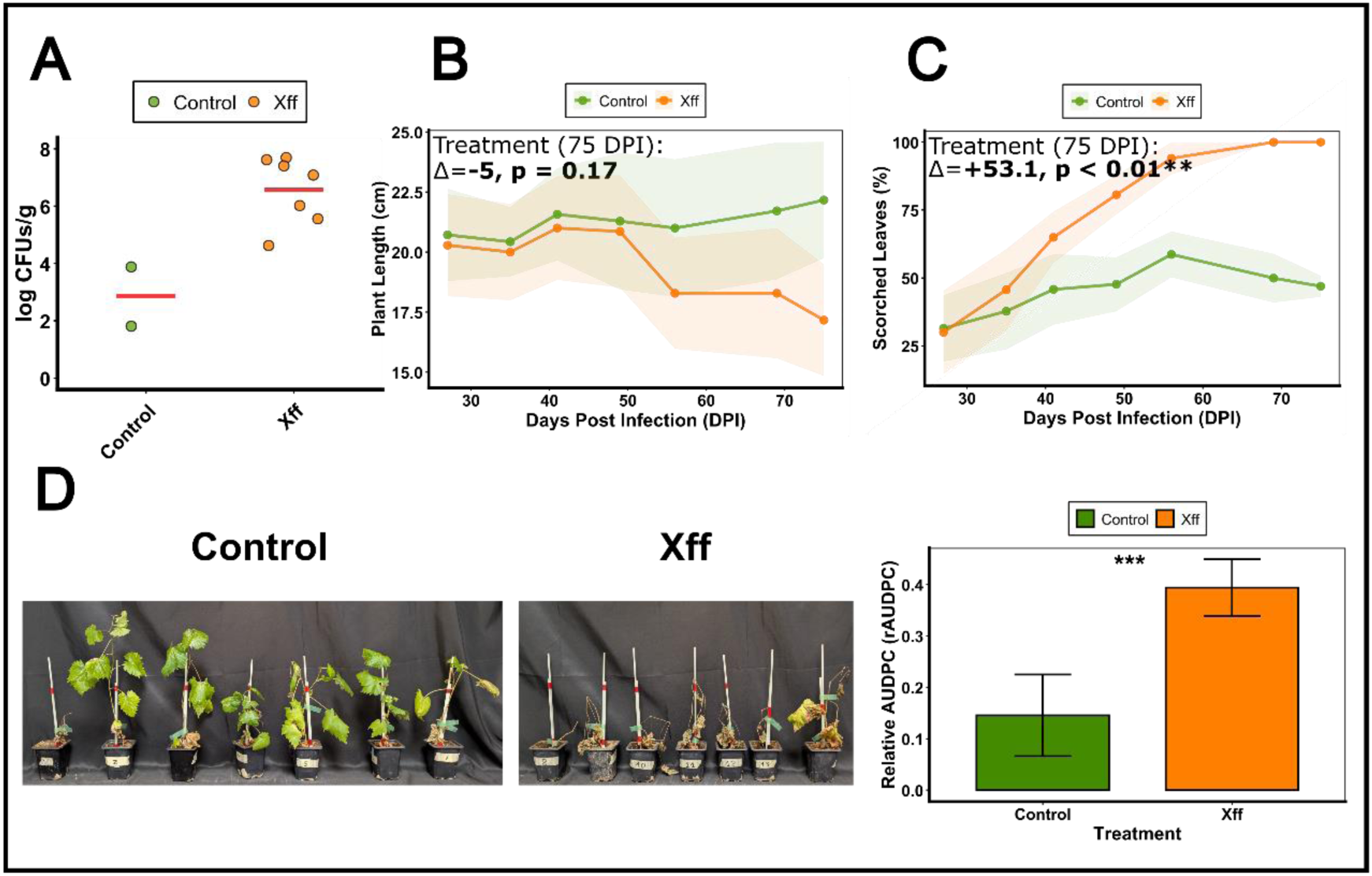
Assessment of *Xylella fastidiosa* colonization and disease progression in *Vitis vinifera*. (A) Estimated bacterial population levels in petiole tissues at 35 DPI. Each point represents an individual plant replicate. Red horizontal lines indicate the mean log (CFU g^-1^) value for each treatment: non-inoculated controls (green), *X. fastidiosa* subsp. *fastidiosa* (Xff, orange). (B) Plant growth dynamics over time following inoculation with Xff (orange) compared to non-inoculated controls (green). Shaded areas indicate SEM. The effects of treatment, DPI, and their interaction were assessed using a two-way ANOVA. (C) Dynamics of leaf scorch development over time. Values represent the percentage of scorched leaves per plant replicate for control (green) and Xff-inoculated (orange) plants. (D) Representative non-inoculated control plants and plants inoculated with Xff at 69 days post-inoculation (DPI) (left), alongside the relative Area Under the Disease Progress Curve (rAUDPC) for symptom severity (right). Symptom scores were recorded throughout the experimental period and normalized to the maximum theoretical disease progress area under the disease progress curve. Bars represent mean ± standard error (SEM); green and orange correspond to control and Xff-inoculated plants, respectively. Pairwise comparison between treatments was performed using estimated marginal means (Tukey-adjusted). Significance codes: p ≤ 0.01; **, p ≤ 0.001; ***.

*V. vinifera* also displayed clear disease development following *X. fastidiosa* inoculation (Fig. 5D). Visual inspections at 63 DPI showed that all control plants, except one, maintained green foliage with negligible senescence. Conversely, the inoculated subjects experienced mortality, displaying widespread necrotic scorching. A single plant escaped death but exhibited severe chlorosis and leaf burn (Fig. 5D). rAUDPC values in inoculated plants reached approximately 0.40, representing a significant increase relative to controls (≈0.14, p < 0.0001), indicating substantial and quantifiable disease progression (Fig. 5D and Table S. 1). **Fig. *5***These values were among the highest recorded across the species evaluated and indicated rapid and reproducible bacterial establishment.

Plant length showed a no significant overall effect of treatment (χ^2^ = 0.36, p = 0.55). However, the treatment x DPI interaction was highly significant (χ^2^= 11.00, p < 0.001), as that growth patterns progressive diverged between treatments after 50 DPI (Fig. 5B and Table S. 2). Whilst individual timepoint comparisons did not reach statistical significance, inoculated plants displayed consistent suppression relative to controls, with an estimated mean difference of 5 cm at 75 DPI (p = 0.17).

Foliar symptomatology differed markedly between treatments (Fig. 5C and Table S. 3), with highly significant treatment effects (χ^2^ = 10.85, p < 0.001), DPI effects (χ^2^ = 65.55, p < 0.001), and treatment x DPI interaction (χ^2^ = 28.83, p < 0.001). The proportion of scorched leaves was significantly higher in inoculated plants from 41 DPI onwards (p = 0.047). At 56 DPI, inoculated plants displayed 97.4% scorched leaves, compared to 61.6% in controls. At the final assessment, conducted at 75 DPI, the proportion of scorched leaves was 53.1 percentage points higher in inoculated plants than in controls (p<0.01).

The Tem1-*V. vinifera* combination provided a highly responsive naturally susceptible woody-host system. The species exhibited rapid and consistent bacterial colonisation, quantifiable disease progression, and clear foliar symptoms, supporting its suitability as a woody host model for investigating *X. fastidiosa* pathogenesis and evaluating potential management strategies under controlled growth-chamber conditions.

### *N. tabacum* and *V. vinifera* show the strongest and most consistent responses to *X. fastidiosa* inoculation

The combined analysis of the four host species revealed marked differences in rAUDPC, stem length and leaf scorch responses to *X. fastidiosa* inoculation (Table S. 1, Table S. 2 and Table S. 3, respectively). For rAUDPC, the combined-species analysis (Table S1A) revealed significant main effects of both treatment (p < 0.001) and species (p < 0.001), as well as a significant treatment x species interaction (p = 0.011). Thus, the effect of Xff inoculation on overall disease progression varied significantly among host species. Together with the species-specific analyses (Table S1B), these results identify *N. tabacum* and *V. vinifera* as the hosts showing the strongest and most consistent combination of *X. fastidiosa* colonization and quantifiable disease development under growth chamber conditions.

For stem elongation, the combined model detected no significant overall effect of Xff inoculation across species (p = 0.295). However, highly significant main effects were detected for species (p < 0.001) and DPI (p < 0.001), together with a significant species-by-DPI interaction (p < 0.001). These results indicate that plant growth dynamics differed markedly among host species. The treatment x species interaction was not significant (p = 0.214), providing no overall statistical evidence that the effect of inoculation on stem elongation differed among species. Nevertheless, separate within-species analyses detected a significant treatment effect only in *N. tabacum* (p < 0.001), whereas *M. sativa*, *N. benthamiana*, and *V. vinifera* showed no significant overall treatment effects.

However, this result does not by itself demonstrate that the treatment effect differed among species, particularly because within-species variability and the relatively small sample size for *V. vinifera* (n = 7) may have limited the power to detect the treatment × species interaction. Overall, stem elongation showed limited sensitivity as a common indicator of infection across host species, highlighting the importance of combining growth measurements with leaf scorch development and bacterial titres.

## Discusion

Geographic and infrastructural constraints severely restrict global access to *X. fastidiosa* research. Quarantine regulations and the need for specialized biosafety facilities limit the experimental evaluation of virulence factors, resistant cultivars, and therapeutic interventions. A rapid, reproducible laboratory-based infection protocol applicable to diverse hosts would facilitate research and accelerate the systematic screening of candidate treatments prior to greenhouse or field validation. In this study, we developed and evaluated a microinjection-based inoculation protocol in four plant species to assess its performance as a standardized, accessible tool for *X. fastidiosa* research.

The protocol showed markedly variable efficacy across the four plant hosts evaluated, revealing substantial variation in susceptibility, bacterial establishment, and symptom development. These results emphasize that host selection is a critical component of the experimental design and determines the suitability of each plant system for preliminary screening or subsequent biological validation.

### Performance and limitations of the growth chamber model

The interpretation and extrapolation of our findings should account for the inherent limitations of controlled growth chambers. These systems differ from natural conditions due to reduced thermal and photoperiodic fluctuations, altered light quality and daily light integral, and internal microclimatic gradients in light intensity and airflow. Root-volume and canopy restrictions, simplified biotic interactions with soil microbiota, and equipment-specific variation may also introduce physiological biases. An additional limitation is that natural *X. fastidiosa* transmission occurs through xylem-feeding insect vectors.

Our microinjection method bypasses this transmission stage, preventing the assessment of vector-mediated selection pressures and transmission efficiency (Poorter *et al*., 2012, 2016; Porter *et al*., 2015; Wang & Chen, 2022; Heuermann *et al*., 2023; Karitter *et al*., 2026). Nevertheless, growth chambers provide a valuable and effective experimental system for the preliminary screening of candidate treatments under controlled conditions prior to field validation.

### Host- and strain-dependent performance of the microinjection protocol

The four plant species differed markedly in bacterial establishment and disease expression. *M. sativa* showed the poorest performance among the hosts evaluated. The absence of disease symptoms, limited and inconsistent bacterial detection, and comparable growth trajectories in inoculated and control plants indicate that this species does not consistently support systemic infection under our experimental conditions. Kubaa et al. (2019) previously reported localized necrosis and limited systemic colonization following needle inoculation with Xfp strain “De Donno”. To account for potential subspecies-dependent differences, we tested this Xfp strain, together with Xff, and Xfm strains in *M. sativa*, but none produced consistent colonization or disease development. We therefore selected Xff for the remaining hosts to standardize the cross-species comparison. Our findings do not exclude the possibility that *M. sativa* supports particular *X. fastidiosa* strains or responds under alternative inoculation conditions, but they indicate that it has limited utility for experiments requiring reproducible systemic infection.

Nicotiana species have been extensively used to investigate *X. fastidiosa* infection and potential treatments, mostly in biosafety greenhouses or affected field regions (Lopes *et al*., 2000, 2020; Alves *et al*., 2003; De La Fuente *et al*., 2013; Gómez *et al*., 2020; Pereira *et al*., 2020; El Handi *et al*., 2022; Baró *et al*., 2022; Shantharaj *et al*., 2023; Incampo *et al*., 2025; Buisac *et al*., 2026). To our knowledge, only two studies have evaluated *X*. *fastidiosa* infection under controlled growth-chamber conditions, both using *N. tabacum* (Francis *et al*., 2008; Tatulli *et al*., 2024).

In our study, the two *Nicotiana* species displayed contrasting levels of experimental reproducibility. *N. benthamiana* inoculated with Xff strain 5235 showed marked growth inhibition, but bacterial detection and establishment were variable. Moreover, four of the six control plants died within the first month, resulting in high background mortality and limiting the interpretation of symptom differences between treatments. Plant deterioration by 29 DPI may also have reduced the reliability of bacterial detection by qPCR. Therefore, *N. benthamiana* did not provide a sufficiently robust or reproducible growth-chamber model under the conditions tested.

By contrast, *N. tabacum* plants inoculated with Xff Tem1 exhibited more reproducible bacterial establishment, sustained bacterial titres of 6.0–8.2 log CFU g⁻¹, and clear phenotypic differences relative to controls within 4-6 weeks. These findings agree with previous reports (Francis et al., 2008; Tatulli et al., 2024), although symptom expression has differed among studies. Although Tatulli et al. (2024) reproduced the inoculation procedure described by Francis et al. (2008) and confirmed bacterial colonization, the authors did not observe the characteristic leaf scorch symptoms reported with the original Francis protocol. Despite this variability in symptom expression across studies, the consistent bacterial establishment observed in *N. tabacum*, together with its manageable growth requirements and relatively rapid response, supports its use for the preliminary screening of antibacterial compounds, virulence factors, and other candidate interventions.

*V. vinifera* provided a relevant woody-host model and supported rapid systemic infection.

All inoculated plants were positive at the first sampling point, 32 DPI, with bacterial titres ranging from 4.6 to 7.7 log CFU g⁻¹. Infection was accompanied by increased rAUDPC values and extensive leaf scorch, whereas stem elongation was not significantly affected. The absence of a detectable effect on plant height may partly reflect canopy restrictions within the growth chamber, which could mask differences in internode elongation. Leaf scorch and bacterial titre therefore appear to be more informative disease indicators than vegetative height in chamber-grown grapevine plants.

Our findings align with those of Feil and Purcell (2001), who showed systemic infection in seed-derived, potted *V. vinifera* plants under growth-chamber conditions. Whereas their study focused primarily on temperature-dependent bacterial growth, the present work comparatively evaluates microinjection across multiple host species. Overall, the rapid colonization observed and the bacterial titres obtained support the use of seed-derived grapevine plants under growth-chamber conditions.

### Biological basis of host-dependent responses

The contrasting responses among the plant species probably reflect interactions between pathogen colonization traits and host immune, anatomical, and physiological characteristics. Bacterial establishment is influenced by the capacity of *X. fastidiosa* to evade initial immune recognition and by the compatibility of its cell wall-degrading enzymes with xylem pit-membrane composition, which may permit or restrict movement between vessels(Luvisi *et al*., 2017; Huang *et al*., 2020; Raffini *et al*., 2020; Castro *et al*., 2021; Landa *et al*., 2022). Vascular architecture, bacterial adhesion to host surfaces, and host–pathogen evolutionary history may also affect bacterial movement, population regulation, and the transition from asymptomatic colonization to disease (Purcell, 2013; Castro *et al*., 2021).

Disease severity may additionally depend on the magnitude and regulation of host vascular responses. Susceptible plants may experience hydraulic dysfunction associated with extensive vessel occlusion by tyloses and gums (Castro *et al*., 2021; Picciotti *et al*., 2021; Serio *et al*., 2024; Mourou *et al*., 2025), whereas resistant plants may restrict occlusion and reinforce vascular tissues through lignification (Saponari *et al*., 2019; Raffini *et al*., 2020; Morelli *et al*., 2021; Serio *et al*., 2024). Differences in the composition and stability of the endophytic microbiome could also influence pathogen proliferation through competitive or antagonistic interactions.(Purcell, 2013; Landa *et al*., 2022; Serio *et al*., 2024; Mourou *et al*., 2025).

These mechanisms provide plausible explanations for the host-dependent responses observed here, particularly the contrast between inconsistent establishment in *M. sativa* and *N. benthamiana* and reproducible systemic infection in *N. tabacum* and *V. vinifera*. Targeted analyses of vascular anatomy, defense activation, bacterial distribution, and microbiome composition will therefore be required to establish the mechanisms responsible for these differences.

### Practical applications and model selection criteria

This work provides a practical framework for selecting plant models according to the experimental objective. *N. tabacum* was the most suitable herbaceous host evaluated for rapid, cost-effective preliminary screening, combining relatively reproducible bacterial establishment with measurable symptom development within 4-6 weeks. Moreover, N. tabacum is itself a crop host affected by X. fastidiosa, particularly in South America, which adds biological relevance to this experimental system (Lopes *et al*., 2020). It may therefore be useful for prioritizing antimicrobial compounds, candidate virulence factors, or therapeutic interventions before conducting more resource-intensive experiments.

*V. vinifera* provides a complementary woody-host system for validating selected candidates and examining infection in an economically relevant host. Although grapevine requires more space and longer cultivation than *N. tabacum*, it better represents the vascular characteristics and disease responses associated with Pierce’s disease. A sequential strategy in which candidates are first screened in *N. tabacum* and subsequently validated in *V. vinifera* could therefore improve experimental efficiency while maintaining biological relevance.

By contrast, *N. benthamiana* may be less appropriate when consistent bacterial establishment is required, although it could remain useful for particular molecular applications. *M. sativa* showed limited and inconsistent establishment and is therefore not recommended under the conditions evaluated here for studies requiring reproducible systemic infection. However, its performance with other bacterial strains, inoculation procedures, or environmental conditions cannot be excluded.

In conclusion, host choice strongly influenced the performance of the microinjection protocol under growth-chamber conditions. N. tabacum provided the most reproducible herbaceous system for preliminary screening, whereas V. vinifera offered a relevant woody-host model for subsequent validation. Their sequential use provides a practical framework for prioritising control strategies, virulence factors, and resistance-related traits before greenhouse or field evaluation.

## Acknowledgments

This work was funded by research grants awarded to M.R.: the MaX- CSIC Excellence Award (DEEP-MaX-2024_IBBTEC) and MCIN/AEI/10.13039/501100011033 (grant PID2024-155420OB-I00). M.R. was supported by PTQ-17-09029 and RYC2022-035122-I (MCIN/AEI). D.L. was supported by the University of La Rioja (FPI-UR/CAR 2022). A. C. was supported by PTA2024-024827-I (MCIN/AEI). We thank Blanca Landa and Laura Montesinos for providing the bacterial strains and *N. benthamiana* seeds, respectively. We are also grateful to all members of the Intergenomics group for their stimulating discussions.

## Data availability statement

All data supporting the findings of this study are included in the article and its Supplementary Data.

---

**Fig. S.1.**
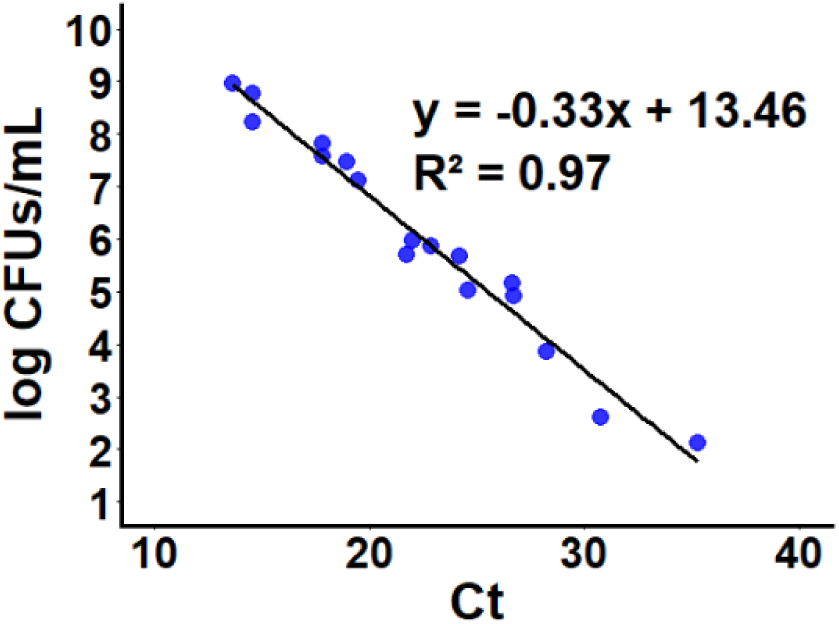
Standard calibration curve for qPCR quantification of *Xylella fastidiosa*. Linear regression of Ct values against log (CFU ml^-1^) using serial dilutions of Xff suspension (OD_600_ = 6) mixed with homogenized plant tissue. The equation log (CFU ml^-1^) = -0.33Ct+13.46 was used to estimate bacterial loads in experimental samples.

## References

1. Alves E, Kitajima EW, Leite B, 2003. Interaction of *Xylella fastidiosa* with Different Cultivars of *Nicotiana tabacum* : a Comparison of Colonization Patterns. Journal of Phytopathology 151, 500–506.

2. Baccari C, Antonova E, Lindow S, 2019. Biological control of Pierce’s disease of grape by an endophytic bacterium. Phytopathology 109, 248–256.

3. Baró A, Montesinos L, Badosa E, Montesinos E, 2021. Aggressiveness of Spanish Isolates of *Xylella fastidiosa* to Almond Plants of Different Cultivars Under Greenhouse Conditions. Phytopathology 111, 1994–2001.

4. Baró A, Mora I, Montesinos L, Montesinos E, 2020. Differential Susceptibility of *Xylella fastidiosa* Strains to Synthetic Bactericidal Peptides. Phytopathology 110, 1018– 1026.

5. Baró A, Saldarelli P, Saponari M, Montesinos E, Montesinos L, 2022. *Nicotiana benthamiana* as a model plant host for *Xylella fastidiosa*: Control of infections by transient expression and endotherapy with a bifunctional peptide. Frontiers in Plant Science 13, 1–12.

6. Bragard C, Dehnen-Schmutz K, Di Serio F et al., 2019. Effectiveness of in planta control measures for *Xylella fastidiosa*. EFSA Journal 17, 1–17.

7. Buisac A, Gascón B, Montesinos E, Montesinos L, 2026. Aggressiveness of Subspecies and Sequence Types of *Xylella fastidiosa* and Plant Response in *Nicotiana benthamiana*. Phytopathology 116, 918–926.

8. Burbank L, Gomez L, Shantharaj D et al., 2024. Virulence Comparison of a Comprehensive Panel of *Xylella fastidiosa* Pierce’s Disease Isolates from California. Plant Disease 108, 1555–1564.

9. Castro C, DiSalvo B, Roper MC, 2021. *Xylella fastidiosa* : A reemerging plant pathogen that threatens crops globally. Plos Pathogens 17, 1–6.

10. Cavalieri V, Fasanelli E, Furnari G et al., 2025. Update of the *Xylella* spp. host plant database – Systematic literature search up to 30 June 2024. EFSA Journal 23, 1–70.

11. Cornara D, Sicard A, Zeilinger AR, Porcelli F, Purcell AH, Almeida RPP, 2016. Transmission of Xylella fastidiosa to Grapevine by the Meadow Spittlebug. Ecology and Epidemiology 106, 1285–1290.

12. Das M, Bhowmick TS, Ahern SJ, Young R, Gonzalez CF, 2015. Control of Pierce’s Disease by phage. PLoS ONE 10, 1–15.

13. Davis MJ, French WJ, Schaad NW, 1981. Isolation Media for the Pierce’s Disease Bacterium. Current Microbiology 6, 309–314.

14. Feil H, Purcell AH, 2001. Temperature-dependent growth and survival of *Xylella fastidiosa* in vitro and in potted grapevines. Plant Disease 85, 1230–1234.

15. Francis M, Civerolo EL, Bruening G, 2008. Improved bioassay of *Xylella fastidiosa* using *Nicotiana tabacum* cultivar SR1. Plant Disease 92, 14–20.

16. Fry WE, 1978. Quantification of General Resistance of Potato Cultivars and Fungicide Effects for Integrated Control of Potato Late Blight. Phytopathology 68, 1650.

17. Ghanbari D, Hasanzadeh N, Ghayeb Zamharir M, Nasr S, El Handi K, Elbeaino T, 2024. Detection and characterization of *Xylella fastidiosa* in Iran: frst report in alfalfa (*Medicago sativa*). Phytopathologia Mediterranea 63, 335–342.

18. Gómez LM, Teixeira-Silva NS, Caserta R, Takita MA, Marques MOM, de Souza AA, 2020. Overexpression of *Citrus reticulata* SAMT in *Nicotiana tabacum* increases MeSA volatilization and decreases *Xylella fastidiosa* symptoms. Planta 252, 1–14.

19. El Handi K, Sabri M, Valentini F et al., 2022. Exploring Active Peptides with Antimicrobial Activity In Planta against *Xylella fastidiosa*. Biology 11, 1–14.

20. Hao L, Johnson K, Cursino L, Mowery P, Burr TJ, 2017. Characterization of the *Xylella fastidiosa* PD1311 gene mutant and its suppression of Pierce’s disease on grapevines. Molecular Plant Pathology 18, 684–694.

21. Harper SJ, Ward LI, Clover GRG, 2010. Development of LAMP and real-time PCR methods for the rapid detection of *Xylella fastidiosa* for quarantine and field applications. Phytopathology 100, 1282–1288.

22. Heuermann MC, Altmann T, Knoch D, 2023. Natural plant growth and development achieved in the IPK PhenoSphere by dynamic environment simulation. Nature Communications 14, 1–14.

23. Hopkins DL, 2005. Biological control of Pierce’s disease in the vineyard with strains of *Xylella fastidiosa* benign to grapevine. Plant Disease 89, 1348–1352.

24. Huang W, Reyes-caldas P, Mann M et al., 2020. Bacterial Vector-Borne Plant Diseases : Unanswered Questions and Future Directions. Molecular Plant 13, 1379–1393.

25. Joint Research Center, 2025. EU quarantine pests: new assessment of the potential impacts.

26. Kahn AK, Sicard A, Cooper ML, Daugherty MP, Donegan MA, Almeida RPP, 2023. Progression of *Xylella fastidiosa* Infection in Grapevines Under Field Conditions. Phytopathology 113, 1465–1473.

27. Karitter P, March-Salas M, Ensslin A et al., 2026. Garden, greenhouse, or climate chamber? Experimental conditions influence whether genetic differences are phenotypically expressed. Plant Biology 2 28, 1–38.

28. Kubaa RA, Giampetruzzi A, Altamura G, Saponari M, 2019. Infections of the *Xylella fastidiosa* subsp. *pauca* Strain “De Donno” in Alfalfa (Medicago sativa) Elicits an Overactive Immune Response. Plants 8, 1–18.

29. Kyrkou I, Pusa T, Ellegaard-Jensen L, Sagot MF, Hansen LH, 2018. Pierce’s disease of grapevines: A review of control strategies and an outline of an epidemiological model. Frontiers in Microbiology 9, 1–23.

30. De La Fuente L, Parker JK, Oliver JE et al., 2013. The Bacterial Pathogen Xylella fastidiosa Affects the Leaf Ionome of Plant Hosts during Infection. PLoS ONE 8, 1– 9.

31. Landa BB, Saponari M, Feitosa-Junior OR et al., 2022. *Xylella fastidiosa*’s relationships: the bacterium, the host plants, and the plant microbiome. New Phytologist 234, 1598–1605.

32. van Leeuwen C, Sgubin G, Bois B et al., 2024. Climate change impacts adaptations of wine production. Nature Reviews Earth and Environment 5, 258–275.

33. Lopes SA, Raiol-j LL, Torres SCZ, Martins EC, Prado SS, Beriam OS, 2020. Differential Responses of Tobacco to the Citrus Variegated Chlorosis and Coffee Stem Atrophy Strains of *Xylella fastidiosa*. Phytopathology 110, 567–573.

34. Lopes SA, Ribeiro DM, Roberto PG, França SC, Santos JM, 2000. *Nicotiana tabacum* as an experimental host for the study of plant-*Xylella fastidiosa* interactions. Plant Disease 84, 827–830.

35. López-Mercadal J, Delgado S, Mercadal P et al., 2021. Collection of data and information in Balearic Islands on biology of vectors and potential vectors of *Xylella fastidiosa* (GP/EFSA/ALPHA/017/01). EFSA Supporting Publications 18, 1–136.

36. López-Mercadal J, Mercadal-Frontera P, Miranda MÁ, 2022. Mechanical management of weeds drops nymphal density of *Xylella fastidiosa* vectors. bioRxiv.

37. Luvisi A, Nicol F, Bellis L De, 2017. Sustainable Management of Plant Quarantine Pests : The Case of Olive Quick Decline Syndrome. Sustainability 9, 1–19.

38. Martínez S, Lacuesta M, Relloso JB, Aragonés A, Herrán A, Ortiz-Barredo A, 2023. European Grapevine Cultivars and Rootstocks Show Differential Resistance to *Xylella fastidiosa* Subsp. fastidiosa. Horticulturae 9, 1–18.

39. Morelli M, García-Madero JM, Jos Á et al., 2021. *Xylella fastidiosa* in olive: A review of control attempts and current management. Microorganisms 9, 1–21.

40. Mourou M, Incampo G, Carlucci M et al., 2025. Insight into biological strategies and main challenges to control the phytopathogenic bacterium *Xylella fastidiosa*. Frontiers in Plant Science 16, 1–13.

41. Nascimento R, Gouran H, Chakraborty S et al., 2016. The Type II Secreted Lipase/Esterase LesA is a Key Virulence Factor Required for *Xylella fastidiosa* Pathogenesis in Grapevines. Scientific Reports 6, 1–17.

42. Pereira WEL, de Andrade SMP, Del Ponte EM et al., 2020. Severity assessment in the *Nicotiana tabacum-Xylella fastidiosa* subsp. *pauca* pathosystem: design and interlaboratory validation of a standard area diagram set. Tropical Plant Pathology 45, 710–722.

43. Perelló A, Romero-Munar A, Martinez SI et al., 2025. Xylem Sap Mycobiota in Grapevine Naturally Infected with *Xylella fastidiosa*: A Case Study: Interaction of *Xylella fastidiosa* with *Sclerotinia sclerotiorum*. Plants 14, 1–16.

44. Picciotti U, Lahbib N, Sefa V, Porcelli F, Garganese F, 2021. Aphrophoridae role in *Xylella fastidiosa* subsp. *pauca* ST53 invasion in southern Italy. Pathogens 10, 1– 25.

45. Poorter H, Fiorani F, Pieruschka R, Putten WH Van Der, Kleyer M, Schurr U, 2016. Tansley review Pampered inside , pestered outsideDifferences and similarities between plants growing in controlled conditions and in the field. New Phytologist 212, 838–855.

46. Poorter H, Fiorani F, Stitt M et al., 2012. The art of growing plants for experimental purposes : a practical guide for the plant biologist. Functional Plant Biology 39, 821–838.

47. Portaccio L, Vergine M, Bene A et al., 2025. Chemical reatments Tested Against *Xylella fastidiosa*: Strategies, Successes and Limitations. Pathogens 14, 1–25.

48. Porter AS, Evans C, Gerald F, Mcelwain JC, Yiotis C, Kingston CE, 2015. How well do you know your growth chambersTesting for chamber effect using plant traits. Plant Methods 11, 1–10.

49. Purcell A, 2013. Paradigms : Examples from the Bacterium *Xylella fastidiosa*. Annual Review of PhytopathologyPhytopathology 51, 339–356.

50. Raffini F, Bertorelle G, Biello R, D’Urso G, Russo D, Bosso L, 2020. From Nucleotides to Satellite Imagery : Approaches to Identify and Manage the Invasive Pathogen *Xylella fastidiosa* and Its Insect Vectors in Europe. Sustainability 12, 1–38.

51. Rodríguez M, 2019. Estudio del origen genético de la variedad de vid Garnacha Blanca , de su diversidad fenotípica y de los efectos moleculares asociados a la variación en el color de la uva. Universidad de La Rioja.

52. Rogers EE, 2012. Evaluation of *Arabidopsis thaliana* as a Model Host for *Xylella fastidiosa*. Molecular Plant-Microbe Interactions 25, 747–754.

53. Saponari M, Boscia D, Altamura G et al., 2017. Isolation and pathogenicity of *Xylella fastidiosa* associated to the olive quick decline syndrome in southern Italy. Scientific Reports 7, 1–13.

54. Saponari M, Giampetruzzi A, Loconsole G, Boscia D, Saldarelli P, 2019. *Xylella fastidiosa* in olive in apulia: Where we stand. Phytopathology 109, 175–186.

55. Serio F, Imbriani G, Girelli CR, Miglietta PP, Scortichini M, Fanizzi FP, 2024. A Decade after the Outbreak of *Xylella fastidiosa* subsp. *pauca* in Apulia (Southern Italy): Methodical Literature Analysis of Research Strategies. Plants 13, 1–54.

56. Shantharaj D, Naranjo E, Merfa M V, Cobine PA, Santra S, Fuente LD La, 2023. Zinc Oxide-Based Nanoformulation Zinkicide Mitigates the Xylem-Limited Pathogen *Xylella fastidiosa* in Tobacco and Southern Highbush Blueberry. Plant Disease 107, 1096–1106.

57. Tatulli G, Baldassarre F, Schiavi D et al., 2024. Chitosan-Coated Fosetyl-Al Nanocrystals’ Efficacy on Nicotiana tabacum Colonized by *Xylella fastidiosa*. Phytopathology 114, 1466–1479.

58. Wallis CM, Gorman Z, 2024. Pre-inoculation water deficit effects on grapevine physiology, *Xylella fastidiosa* titers, and Pierce’s disease progression. BMC Research Notes 17, 1–6.

59. Wang J, Chen C, 2022. From Laboratory to Field : The Effect of Controlling Oscillations in Temperature on the Growth of Crops. Horticulture 8, 1–14.

60. Yang L, Lin H, Takahashi Y, Chen F, Walker MA, Civerolo EL, 2011. Physiological and Molecular Plant Pathology Proteomic analysis of grapevine stem in response to *Xylella fastidiosa* inoculation. Physiological and Molecular Plant Pathology 75, 90– 99.

61. Zhang S, Jain M, Fleites LA, Rayside PA, Gabriel DW, 2019. Identification and characterization of menadione and benzethonium chloride as potential treatments of Pierce’s disease of grapevines. Phytopathology 109, 233–239.

