## Supplemental figures 1 and 2 and supplemental tables 1-3 for "*Nicotiana tabacum* and *Vitis vinifera* as suitable growth-chamber models for microinjection-based studies of *Xylella fastidiosa*"

### A microinjection-based infection system for *Xylella fastidiosa* enabling reproducible multi-host assays in laboratory growth chambers

David Labarga[1,2*], Ana Cuevas[2], Miguel López-Belmonte[2], Natalia García-Tomsig[2], and Marta Robledo[2*]

*1.- Instituto de Ciencias de la Vid y del Vino (ICVV), CSIC - Gobierno de la Rioja - Universidad de La Rioja, Logroño 26007, Spain.*

*2.- Instituto de Biomedicina y Biotecnología de Cantabria (IBBTEC), Universidad de Cantabria-Consejo Superior de Investigaciones Científicas (CSIC), Santander, Spain*

#### Supplementary Material

**Fig. S. 1.** Standard calibration curve for qPCR quantification of *Xylella fastidiosa*.

**Fig. S. 2.** Pictures at 213 days post-inoculation of non-inoculated control plants, plants infected with *X. fastidiosa* subsp. *multiplex* (Xfm) and *X. fastidiosa* subsp. *pauca* (Xfp).

**Table S. 1.** GLMM statistical results of rAUDPC following *Xylella fastidiosa* inoculation in *Medicago sativa, Nicotiana benthamiana, Nicotiana tabacum* and *Vitis vinifera.*

**Table S. 2.** GLMM statistical results of stem length following *Xylella fastidiosa* inoculation in *Medicago sativa, Nicotiana benthamiana, Nicotiana tabacum* and *Vitis vinifera..*

**Table S. 3.** Leaf scorch statistical results in *Nicotiana benthamiana*, *Nicotiana tabacum*, and *Vitis vinifera* inoculated with *Xylella fastidiosa*.

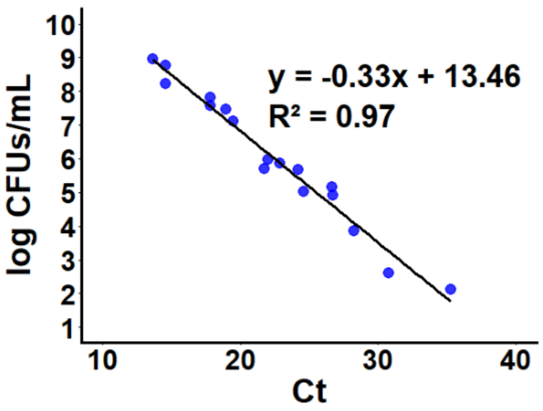


**Fig. S. 1. Standard calibration curve for qPCR quantification of *Xylella fastidiosa*.** Linear regression of Ct values against log (CFU $ml^{-1}$) across serial dilutions of bacterial suspension ($OD_{600}$ = 6) mixed with plant tissue. The equation log (CFU $ml^{-1}$) = -0.33Ct+13.46 enables conversion of Ct values to bacterial load.) = -0.33Ct+13.46 enables conversion of Ct values to bacterial load.

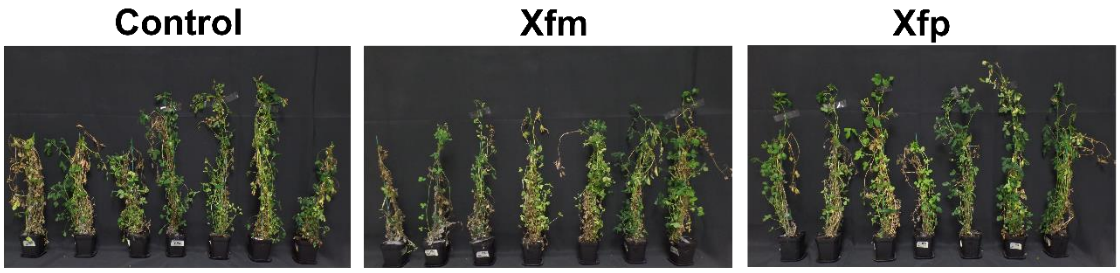


**Fig. S. 2.** Pictures at 213 days post-inoculation of non-inoculated control plants, plants infected with *X. fastidiosa* subsp. *multiplex* (Xfm) and *X. fastidiosa* subsp. *pauca* (Xfp).

**Table S. 1. GLMM statistical results of rAUDPC following *Xylella fastidiosa* inoculation.** (A) Combined-species analysis assessing treatment (T), species (S), and T x S interaction effects on relative Area Under the Disease Progress Curve (rAUDPC). (B) Tukey-adjusted pairwise comparisons between control and Xff-inoculated plants within species. Plant replicate was used as a random effect. Degrees of freedom (df), $\chi 2$ statistics, p-values, and significance levels are reported. Significance codes: $p > 0.05$; * $p \leq 0.05$; *** $p \leq 0.001$.Significance codes: $p > 0.05$; * $p \leq 0.05$; *** $p \leq 0.001$.

A

rAUDPC combined-species

| Factor | df | Chi-square | p-value | Significance |
|---|---|---|---|---|
| Treatment | 1 | 19.816 | 8.53E-06 | *** |
| Species | 3 | 31.77 | 5.85E-07 | *** |
| T x S | 3 | 11.204 | 0.01067 | * |

B

rAUDPC species-specific

| | p-value Post-hoc test |
|---|---|
| *Medicago sativa* | 0.9229 |
| *Nicotiana benthamiana* | 0.9918 |
| *Nicotiana tabacum* | <0.0001*** |
| *Vitis vinifera* | <0.0001*** |

**Table S. 2. GLMM statistical results of stem length following *Xylella fastidiosa* inoculation.** (A) Combined-species analysis evaluating the effects of treatment (T), species (S), days post-inoculation (DPI), and their interactions. (B) Species-specific models assessing treatment and DPI effects. Plant replicate was included as a random effect in all models. Degrees of freedom (df), type II Wald $\chi 2$ statistics, p-values, and significance levels are shown. N.s., $p > 0.05$; * $p \leq 0.05$; ** $p \leq 0.01$; *** $p \leq 0.001$.$\chi 2$ statistics, p-values, and significance levels are shown. N.s., $p > 0.05$; * $p \leq 0.05$; ** $p \leq 0.01$; *** $p \leq 0.001$.

A

Plant length combined-species

| Factor | df | Chi-square | p-value | Significance |
|---|---|---|---|---|
| Treatment | 2 | 2.4403 | 0.29518 | n.s. |
| Species | 5 | 276.728 | <2.2e-16 | *** |
| DPI | 4 | 1098.92 | <2.2e-16 | *** |
| T x S | 3 | 4.4846 | 2.14E-01 | n.s. |
| T x DPI | 3 | 7.0699 | 0.0697 | n.s. |
| S x DPI | 9 | 153.099 | <2.2e-16 | *** |
| T x S x DPI | 9 | 14.955 | 0.09218 | n.s. |

B

Plant length species-specific

| | Factor | Treatment | DPI | T x DPI |
|---|---|---|---|---|
| *Medicago sativa* | df | 1 | 3 | 3 |
| | Chi-square | 0.3805 | 787.574 | 10.2666 |
| | p-value | 0.53735 | <2e-16 | 0.01643 |
| | Significance | n.s. | *** | * |
| *Nicotiana benthamiana* | df | 1 | 3 | 3 |
| | Chi-square | 2.4136 | 17.486 | 12.7725 |
| | p-value | 0.1202854 | 0.00056 | 0.00516 |
| | Significance | n.s. | *** | ** |
| *Nicotiana tabacum* | df | 1 | 3 | 3 |
| | Chi-square | 37.036 | 32.246 | 37.819 |
| | p-value | 1.16E-09 | 4.65E-07 | 3.09E-08 |
| | Significance | *** | *** | *** |
| *Vitis vinifera* | df | 1 | 1 | 1 |
| | Chi-square | 0.3619 | 6.9996 | 11.003 |
| | p-value | 0.5474539 | 0.00815 | 0.00091 |
| | Significance | n.s. | ** | *** |

**Table S. 3. Leaf scorch statistical results in *Nicotiana benthamiana, Nicotiana tabacum*, and *Vitis vinifera* inoculated with *Xylella fastidiosa*.** Treatment effects across days post-inoculation (DPI) were assessed using Fisher's exact test for *N. benthamiana*, and GLMMs (with treatment, DPI, and their interaction as fixed effects, and plant replicate as a random effect) for *N. tabacum* and *V. vinifera*. Type II Wald $\chi 2$ tests statistics, degrees of freedom (df), p-values, and significance levels are shown. Significance codes: ns, $p > 0.05$; * $p \leq 0.05$; ** $p \leq 0.01$; *** $p \leq 0.001$.

| | DPI | p-value |
|---|---|---|
| *Nicotiana benthamiana* | 29 | 0.0272 |

| | Factor | Treatment | DPI | T x DPI |
|---|---|---|---|---|
| *Nicotiana tabacum* | df | 1 | 3 | 3 |
| | Chi-square | 39.133 | 19.694 | 27.874 |
| | p-value | 3.96E-10 | 1.96E-04 | 3.86E-06 |
| | Significance | *** | *** | *** |
| *Vitis vinifera* | df | 1 | 4 | 4 |
| | Chi-square | 10.852 | 65.552 | 28.833 |
| | p-value | 0.0009868 | 3.33E-12 | 6.54E-05 |
| | Significance | *** | *** | *** |
